# KGF induces podosome formation via integrin-Erk1/2 signaling in human immortalized oral epithelial cells

**DOI:** 10.1101/508416

**Authors:** Guoliang Sa, Zhikang Liu, Jiangang Ren, Qilong Wan, Xuepeng Xiong, Zili Yu, Heng Chen, Yifang Zhao, Sangang He

## Abstract

Our recent study established the role of integrins in KGF-induced oral epithelial adhesion and rete peg elongation. However, how extracellular matrix remodeling cooperates with the increased epithelial adhesion during rete peg elongation has yet to be determined. Podosomes are cell-matrix contact structures that combine several abilities, including adhesion and matrix degradation. In the present study, we identified podosome formation at the ventral side of human immortalized oral epithelial cells (HIOECs) upon KGF treatment. Moreover, podosomal components colocalized with the F-actin-cortactin complex and matrix degradation assays demonstrated the ability of the F-actin-cortactin complex to degrade matrix. Inhibition both of integrin subunits β4 and β1 with specific blocking antibodies and inhibition of Erk1/2 abrogated the KGF-induced podosome formation. Notably, knockdown of integrin subunits β4 and β1 with specific siRNA downregulated the phosphorylation levels of Erk1/2. In contrast, inhibition of both Erk1/2 could upregulate the expression of integrin subunits β4 and β1. These results demonstrate that KGF induces podosome formation via integrin-Erk1/2 signaling in HIOECs, suggesting a novel mechanism by which integrins enhance oral epithelial adhesion and rete peg elongation.

**Summary Statement:** Human immortalized oral epithelial cells can form podosomes after stimulation with KGF during which integrin-Erk1/2 signaling play essential roles.

## Introduction

Oral mucosal rete pegs are epithelial protrusions toward the lamina propria formed by epithelial cell proliferation and migration upon various factors such as mechanical stimuli (Lawlor and Kaur 2015; Wu et al. 2013). An increasing number of studies demonstrate that rete pegs contribute to epithelial adhesion because they can enlarge the contact area between the oral epithelium and lamina propria (Clement et al. 2013). However, the extracellular matrices underlying the basement membrane dynamically remodel during the progress of rete peg elongation, which may compromise epithelial adhesion. Consequently, how extracellular matrix (ECM) remodeling cooperates with increased epithelial adhesion during rete peg elongation has yet to be determined.

Podosomes are dynamic cell-matrix contact structures that combine several key abilities, including adhesion and matrix degradation, via their expression of integrins and matrix metalloproteases (MMPs) (Murphy and Courtneidge 2011; Schachtner et al. 2013). Although the first description of podosomes was in Rous sarcoma virus-transformed fibroblasts as aberrant adhesion structures and subsequently in several cell types of mesenchymal origin including osteoclasts and macrophages, they are not unique to cells of mesenchymal origin (Schachtner et al. 2013). Endothelial cells can form podosomes quickly after short-term treatment with TGFβ, TNF-α, VEGF, and phorbol ester (Daubon et al. 2016; Tatin et al. 2006; Varon et al. 2006). Moreover, these podosomes could organize into rosette structures that were the precursors of new vascular branching points (Daubon et al. 2016; Seano et al. 2014). Phorbol esters can also activate human bronchial epithelial cells and have been implicated in not only podosome assembly but also recruitment of MMPs to podosomes for matrix degradation (Xiao et al. 2009). Most importantly, the skin-derived keratinocyte cell line HaCat also displayed podosome-like structures (Spinardi et al. 2004). Based on these studies, we hypothesize that oral epithelial cells have the possibility to form podosomes upon growth factor treatment.

Our previous study demonstrated that KGF simultaneously enhanced oral rete peg elongation and epithelial adhesion, suggesting the possibility that KGF may induce podosomes formation (Sa et al. 2017). In this study, we identified the formation of podosome-like structures (colocalization of F-actin with cortactin) in HIOECs 24 h after KGF treatment by confocal microscopy. Moreover, we analyzed the colocalization of podosomal components including integrins and MMPs with F-actin-cortactin and the ability to degrade matrix. Additionally, we explored the related signaling molecules that regulate podosomes formation in HIOECs after KGF treatment.

## Results

### KGF induces the assembly of podosome structures in HIOECs

To verify our hypothesis that KGF can promote the formation of podosomes, we detected the colocalization of F-actin and cortactin 24 h after KGF treatment by confocal microscopy. As shown in Figure 1, colocalization of F-actin and cortactin was almost invisible in HIOECs of control groups (1A), whereas 40%-50% of HIOECs showed colocalization of F-actin and cortactin 24 h after KGF treatment (1B). These podosome-like structures presented as dot-like structures with a diameter approximately 0.4 μm (Fig. 1 A, magnification). Next, we investigated the podosomal components in the podosome-like structures (identified by colocalization of F-actin-cortactin at the ventral side of the cell). The results showed that integrin subunits α6, β4, α3, β1, and MMP14 colocalized with F-actin in podosome-like structures (Fig. 1C and Supplementary Fig. 1). To ascertain the ability of the podosome-like structures to degrade matrix, we then performed matrix degradation assays. The results demonstrated that gelatin degradation, which has previously been associated with podosomes, was observed in cells with podosomes (Fig. 1D and E).

**Figure 1.**
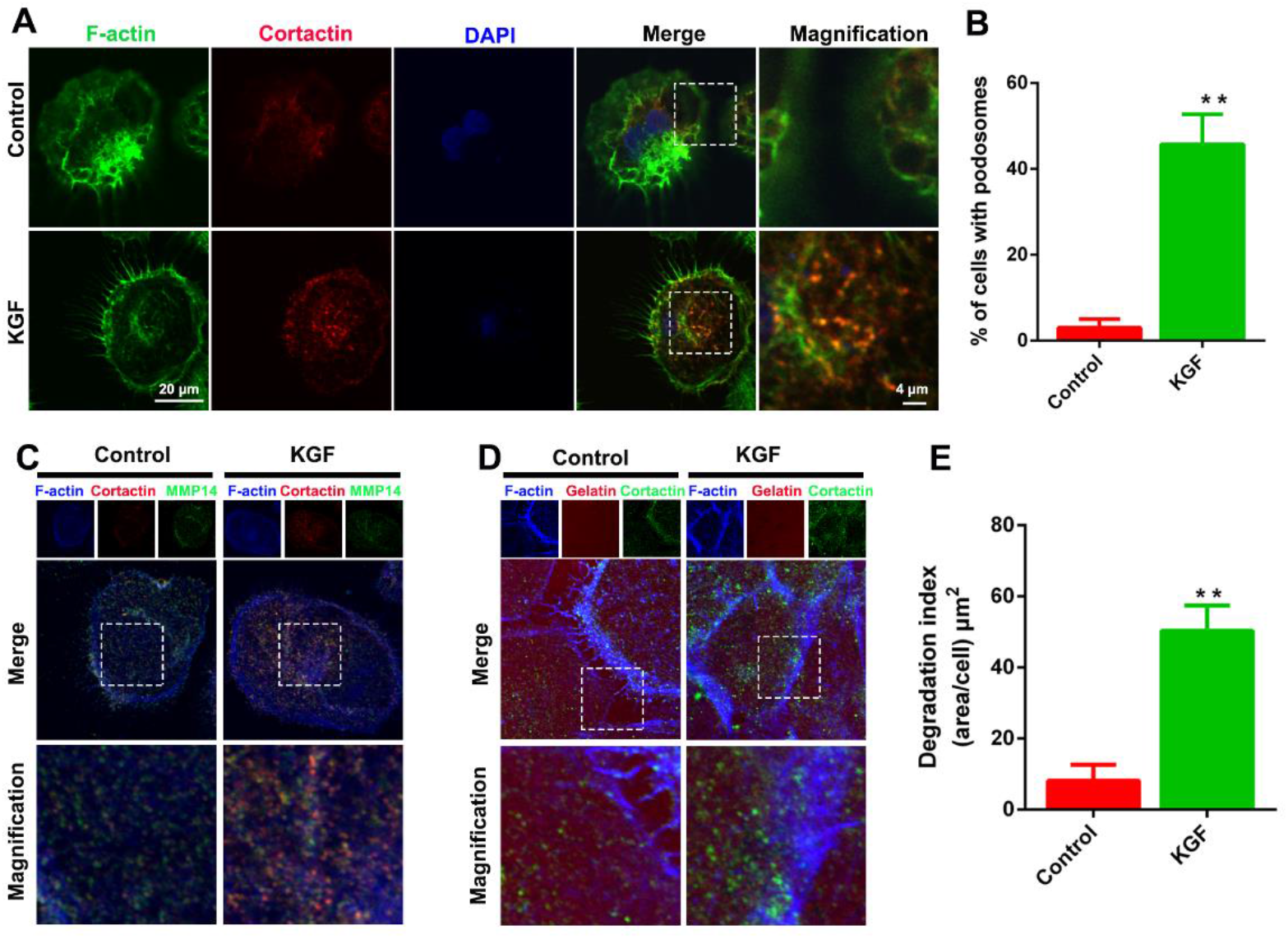
KGF Induces the Assembly of Podosome Structures in HIOECs. (A and B) 24 h after KGF treatment, the ratio of cells containing F-actin-cortactin complexes in the KGF group was significantly elevated compared to the control group. n=3 independent experiments in which 100 cells per experimental point were analyzed. (C) MMP14 was colocalized with F-actin-cortactin complexes in HIOECs of the KGF group. (D, E) The gelatin degradation ability of HIOECs in the KGF group was much higher than the control group. n=3 independent experiments in which 10 cells per experimental point were analyzed. Data are expressed as the means ± SEM. \*\**P* < 0.01, \*\*\**P*< 0.001.

### Integrins are involved in KGF-induced podosome formation

A recent study demonstrated that KGF simultaneously enhanced oral rete peg elongation and epithelial adhesion (Sa et al. 2017). However, inhibition of integrins with HYD-1, an antagonist to integrin α6, β4, α3, and β1, drastically abrogated the KGF-induced oral epithelial adhesion and rete peg elongation, indicating the role of integrins in adhesion and matrix remodeling. Because adhesion and matrix degradation are two major podosome functions, we wonder whether integrins are functionally implicated in KGF-induced podosome formation in HIOECs. Therefore, we pretreated HIOECs with HYD-1 to analyze the impact of integrins on KGF-induced podosome formation. As shown in Fig. 2A and B, HYD-1 significantly abrogated the KGF-induced podosome formation. Importantly, HYD-1 can simultaneously inhibit the function of integrin subunits α6, β4, α3, and β1 (DeRoock et al. 2001). To further analyze the relative importance of integrin subunits α6, β4, α3, and β1 in KGF-induced podosome formation, we treated HIOECs with their specific function-blocking antibodies GoH3, 3E1, P1B5, and 12G10 respectively. As shown in Fig. 2B and Supplementary Fig. 2, inhibition of integrin subunit β4 with 3E1 and β1 with 12G10 significantly impaired podosome formation, while inhibition of integrin subunit α6 with GoH3 and α3 with P1B5 had only a marginal effect on KGF-induced podosome formation.

**Figure 2.**
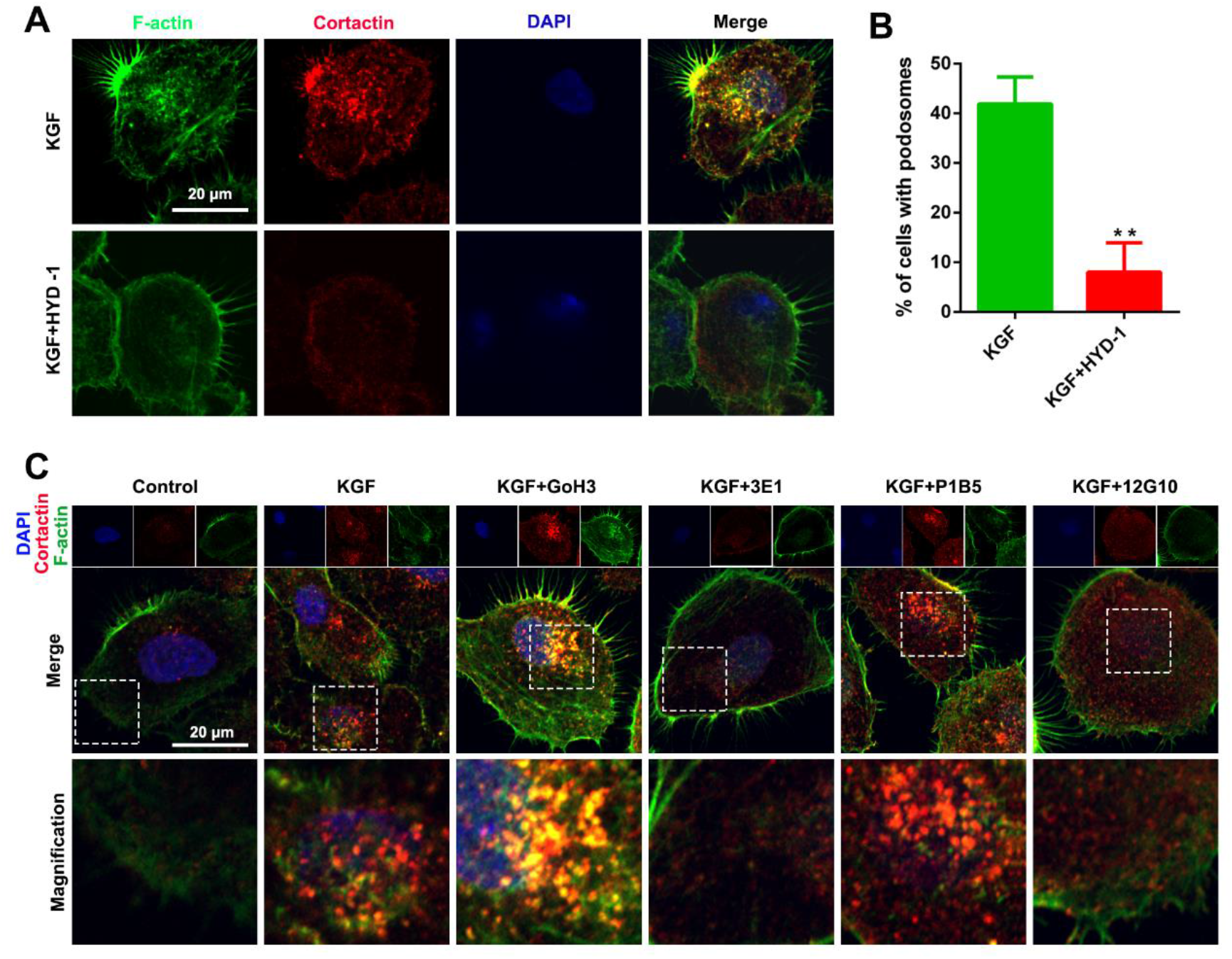
Integrins are involved in KGF-induced podosome formation. (A) HYD-1, an antagonist to integrin subunits α6, β4, α3, and β1, significantly inhibited the KGF-induced podosome formation in HIOECs. (B) n=3 independent experiments in which 100 cells per experiment were analyzed. Data are expressed as the means ± SEM. \*\**P* < 0.01. (C) The impact of function-blocking antibodies against integrin subunits α6 (GoH3), β4 (3E1), α3 (P1B5), and β1 (12G10) on the KGF-induced podosome formation in HIOECs. The statistical analysis is shown in the supplementary.

### KGF mediated its effects on HIOECs through Erk1/2 and Akt signaling

The mitogen-activated protein kinases (MAPK) pathway, the Akt pathway, and the Janus kinase-signal transducers and activators of transcription (Jak-Stat) pathway are three main signaling pathways that mediate the effects of growth factors and cytokines on epithelial cells (Bai et al. 2017; Braun et al. 2006; Canady et al. 2013; Liang et al. 1998). In the present study, we detected the phosphorylation level of Erk1/2, p38, Jnk, Akt, Jak2, and Stat3 in HIOECs by Western blot 15 min after KGF treatment. The results showed that the expression level of KGFR and phosphorylation levels of Erk1/2, Akt, and Stat3 were upregulated upon 10 ng/mL KGF treatment, whereas the phosphorylation levels of p38 and Jnk were unaffected (Fig. 3A-F). Notably, the phosphorylation level of Jak2 was significantly downregulated. Jak2 is considered to be the upstream molecule of Stat3, but their phosphorylation levels showed an inverse relationship. These results indicated that KGF mediated its effects on HIOECs possibly through Erk1/2 and Akt signaling. Because Erk1/2 and Akt play essential roles in cell proliferation, we further analyzed the effects of Erk1/2 and Akt in HIOEC proliferation by EdU and MTT assays. The results demonstrated that U0126 and Ly294002 significantly inhibited the proliferation ability of HIOECs (Fig. 3G-I).

**Figure 3.**
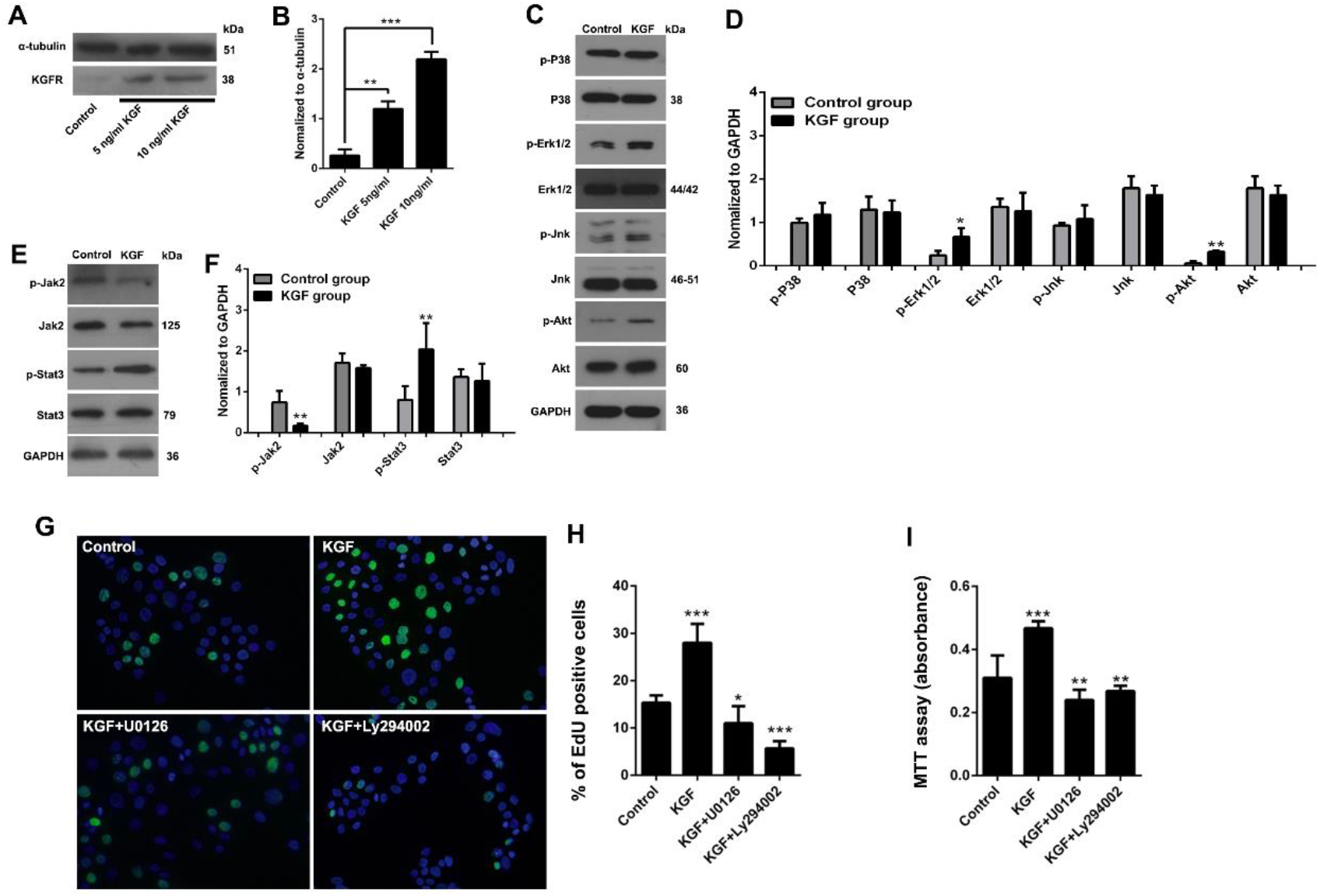
KGF Mediated its Effects on HIOECs through Erk1/2 and Akt Signaling. (A and B) Western blot detection of the expression level of KGFR in HIOECs after KGF treatment, (C and D) the phosphorylation level of Jak2 and Stat3 after KGF stimulation for 15 min, and (E and F) the phosphorylation level of Erk1/2, p38, Jnk, and Akt after KGF stimulation for 15 min. n=3 independent experiments. Analysis of matrix protein expression in protein extracts from each group by ImageJ. (G) EdU labeling assay: the impact of U0126 and Ly294002 on the proliferation ability of HIOECs. (H) The quantification shown in the right graph indicates the mean ± SEM. (I) MTT assay: the impact of U0126 and Ly294002 on the proliferation ability of HIOECs. Data are expressed as the means ± SEM. \**P* < 0.05, \*\**P* < 0.01, \*\*\**P* < 0.001.

### Erk1/2 plays an essential role in KGF-induced podosome formation

Although both Erk1/2 and Akt were significantly activated in HIOECs, we wondered whether both Erk1/2 and Akt were indispensable for KGF-induced podosome formation. Thus, we explored the impact of Erk1/2 and Akt blockade on KGF-induced podosome formation using their specific inhibitors U0126 and Ly294002, respectively. Notably, only inhibition of Erk1/2 abrogated the KGF-induced podosome-like structure formation (Fig. 4A-C and E), whereas inhibition of Akt only had marginal effects on podosome formation in HIOECs (Fig. 4A-B, and D-E).

**Figure 4.**
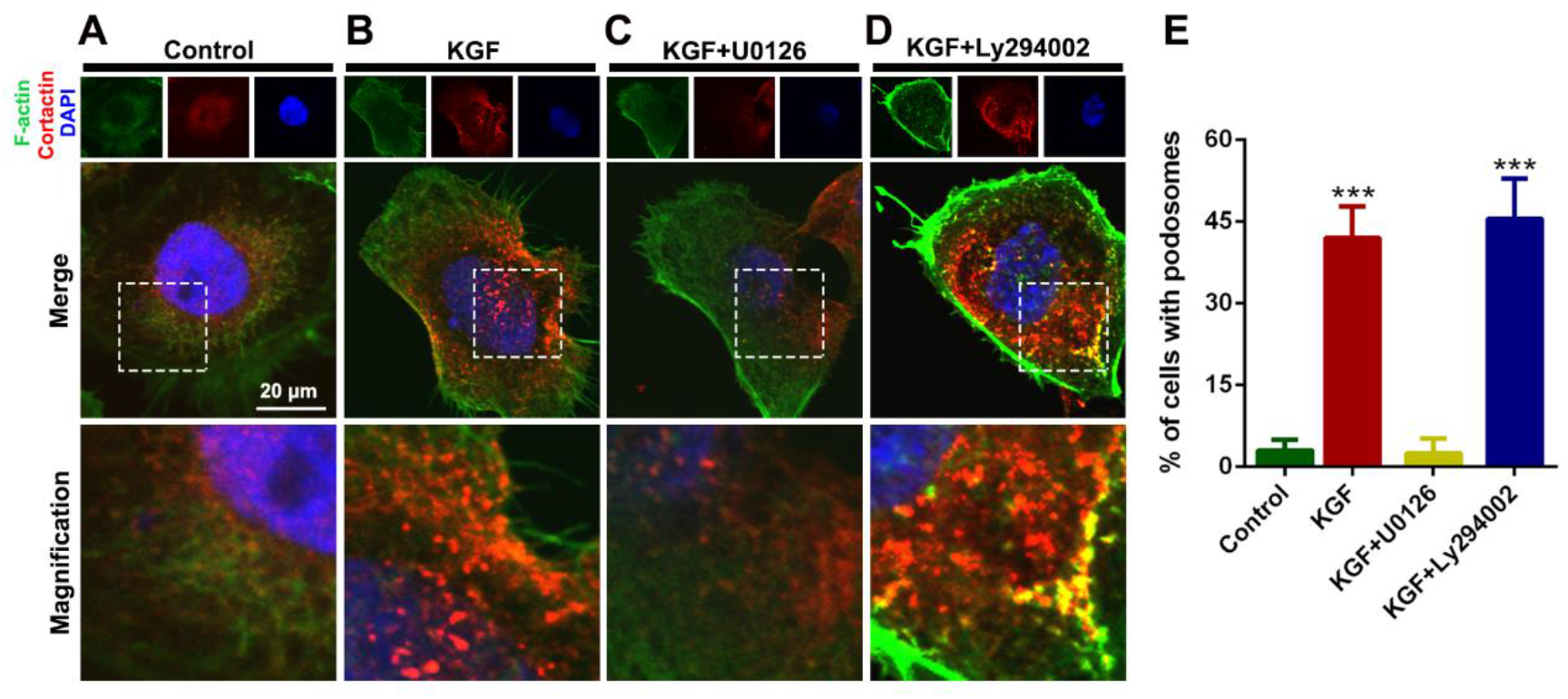
Impact of Erk1/2 and Akt blockade on KGF-induced podosome formation. U0126, specific inhibitor of Erk1/2. Ly294002, specific inhibitor of Akt. (A, B and E) 24 h after KGF treatment, the ratio of cells containing F-actin-cortactin complexes in the KGF group was significantly elevated compared to the control group. (C and E) U0126 significantly inhibited the KGF-induced podosome formation in HIOECs. (D and E) Ly294002 had a marginal effect on KGF-induced podosome formation in HIOECs. n=3 independent experiments in which 100 cells per experiments were analyzed. Data are expressed as the means ± SEM. \*\*\**P*<0.001.

### Integrins are key Mediators between KGF and Erk1/2 activation

As inhibition of integrin subunits β4 and β1 and Erk1/2 abrogated KGF-induced podosome formation, we wondered whether the KGF induced-Erk1/2 activation was associated with integrin subunits β4 and β1 . To test this hypothesis, we suppressed the expression of β4 and β1 integrin with specific siRNAs, verifying the efficiency of the siRNAs by Western blot and real-time PCR (Supplementary Fig. 3A-D). The results showed that knockdown of integrin subunits β4 and β1 significantly downregulated the phosphorylation level of Erk1/2 and Akt in HIOECs (Fig. 5A-D). We further analyzed the impact of specific inhibitors U0126 and Ly294002 on the expression levels of integrins. The results demonstrated that inhibition of both Erk1/2 and Akt significantly upregulated the expression levels of integrin subunits β4 and β1 (Fig. 5E-H). These data suggested that integrin subunits β4 and β1 were essential mediators between KGF and phosphorylation of Erk1/2 and Akt in HIOECs.

**Figure 5.**
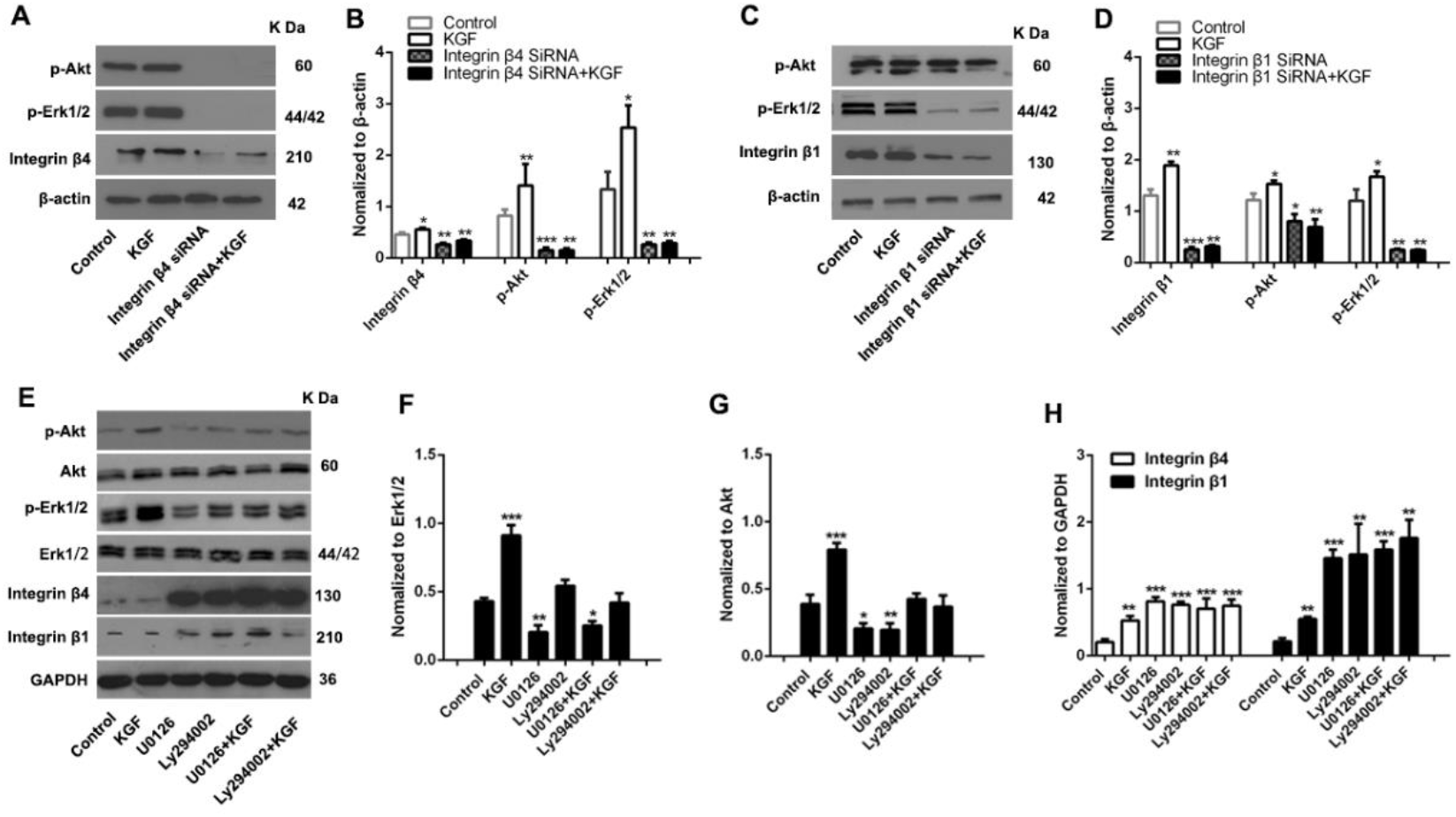
Integrins are Key Mediators between KGF and Activation of Erk1/2. (A and B) Knockdown of integrin β4 by specific siRNA significantly inhibited the expression level of integrin β4 and downregulated the phosphorylation levels of Erk1/2 and Akt. (C and D) Knockdown of integrin β1 by specific siRNA significantly inhibited the expression level of integrin β1 and downregulated the phosphorylation levels of Erk1/2 and Akt. (E-G) U0126 and Ly294002 significantly downregulated the phosphorylation levels of Erk1/2 and Akt. (E and H) U0126 and Ly294002 significantly inhibited the expression levels of integrin subunits β4 and β1. n=3 independent experiments. Data are expressed as the means ± SEM. \**P* < 0.05, \*\**P* < 0.01, \*\*\**P* < 0.001.

## Discussion

In the present study, we explored the possibility of podosome formation in HIOECs after KGF treatment. The results demonstrated that the podosome-like structures in HIOECs presented as dot-like structures with a diameter of 0.4 μm, which were similar in shape to those found in osteoclasts (Linder 2007). Moreover, the key components of podosomes including integrins and MMP14 closely surrounded the F-Actin-cortactin core, suggesting the adhesion and matrix degradation functions of the podosome-like structures. Most importantly, the podosome-like structures in HIOECs indeed had the ability to degrade matrix. Based on these results, we consider that KGF can promote the formation of podosomes in HIOECs, which may be a reasonable mechanism underlying dynamic adhesion during oral mucosal rete peg elongation.

In addition to growth factors and cytokines, integrins, one of the key components of podosomes, also play essential roles in the assembly and function of podosomes in various cell types. For example, β2 integrin participates in podosome formation in dendritic cells, whereas β3 integrin contributes to podosome formation in osteoclasts (Gawden-Bone et al. 2014). A recent study demonstrates that α6β1 integrin is essential for VEGF-induced endothelial podosome rosettes (Seano et al. 2014). The integrin subunits of α6, β4, α3, and β1 are the dominant types expressed in keratinocytes (Margadant et al. 2010). In the present study, we found that the number of podosomes was significantly reduced upon inhibition of the function of integrin subunits β1 and β4, indicating the role of integrin subunits β4 and β1 in podosome assembly in HIOECs. Many studies have demonstrated that integrins also play important roles in podosome self-renewal. For example, Gawden-Bone et al. (2014) found that mutation in amino acids 745 or 746 in integrin β2 inhibited the disassembly of podosomes mediated via Toll-like receptors. Therefore, we hypothesize that integrin subunits β4 and β1 may have a role in different stages of podosome formation and renewal.

In the present study, we found that inhibition of integrins significantly downregulated the KGF-induced phosphorylation of Erk1/2 and Akt, whereas inhibition of Erk1/2 and Akt significantly upregulated the expression of integrin α6, β4, α3, and β1. Therefore, integrins are key mediators between KGF and activation of Erk1/2 and Akt in HIOECs, which is reminiscent of previously described crosstalk between growth factors and integrins (Eliceiri 2001). Integrin ligation has been demonstrated to induce a series of intracellular signaling pathways including MAPK, FAK, Src, small GTPases, and PI3K. Interestingly, these signaling effectors are also activated after growth factor stimulation. Furthermore, integrins can directly associate with growth factor receptors. For example, integrin αvβ3 is directly linked with PDGFR and VEGFR in endothelial cells (Borges et al. 2000; Schneller et al. 1997). More importantly, some growth factors such as IGF interact directly with integrins (Takada et al. 2017). Therefore, the crosstalk between integrins and growth factors and/or growth factor receptors can propagate downstream signaling by regulating the capacity of integrin/growth factor receptor complexes. Presumably, Erk1/2 and Akt may be one of the points of crosstalk between KGF and integrins; however, the concrete mechanisms warrant further investigation.

Our results demonstrated that only Erk1/2 and not Akt plays an indispensable role in KGF-induced podosome formation, which is in accordance with Gu’s study (Gu et al. 2007). However, Akt has also been reported to contribute to the assembly of podosomes in multiple cell types including osteoclasts and endothelial cells (Fey et al. 2016; Kung et al. 2012; Shinohara et al. 2012). Various reasons can explain the diversity of regulatory mechanisms for podosome formation. 1) Both adhesion signaling and growth factors can induce podosome assembly (Hoshino et al. 2013). For example, adhesion signaling is crucial for podosome organization and maturation in osteoclasts, whereas a multiplicity of growth factors including PDGF, VEGF, and TGF-β induce the assembly of podosomes in nonmyeloid cells, such as endothelial cells and smooth muscle cells (Quintavalle et al. 2010). 2) Although podosomes in various cell types show parallel functions in promoting adhesion and matrix remodeling, their morphologies are different (Xiao et al. 2009). For example, in v-Src-transformed NIH 3T3 fibroblasts, podosomes present as rosette-like structures (Seals et al. 2005), whereas phorbol ester-induced podosomes in human airway epithelial cells present as numerous small dots with occasional rosette rings and belts (Xiao et al. 2009). 3) Multiple integrins including integrin subunits β2, β3, α6β1 are involved in the assembly, mutation, and renewal of podosomes in several cell types (Murphy and Courtneidge 2011). Therefore, the downstream signaling pathways mediating these upstream inputs are varied. Notably, downstream signaling pathways involved in podosome assembly always converge on common signaling hubs, especially Rho family GTPases including Rac/Cdc42 and p21-activated kinase (Pak), which ultimately control podosome formation and maturation via regulating cytoskeleton rearrangement (Albiges-Rizo et al. 2009; Hoshino et al. 2013; Linder and Aepfelbacher 2003).

In summary, this work identified podosome assembly in HIOECs after KGF treatment, during which integrin-Erk1/2 signaling plays an indispensable role. The availability of integrins and MMPs within podosomes facilitates the dynamic adhesion and ECM remodeling ability of HIOECs, indicating a possible mechanism by which integrins enhance oral epithelial adhesion and rete peg elongation.

## Materials and Methods

### Antibodies and reagents

Antibodies used for functional blockade are shown in Supplementary Table 1. Antibodies used for Western blot are shown in Supplementary Table 2. The source of HYD-1 (Lys-Ile-Lys-Met-Val-Ile-Ser-Trp-Lys-Gly) and KGF have been described in previous study (Sa et al. 2017). A FITC Phalloidin and an Acti-stain^TM^ 670 phalloidin were obtained from AAT Bioquest Inc. An Erk1/2 inhibitor U0126 and an Akt inhibitor Ly294002 were obtained from Selleck chemicals (Selleck, USA). MTT (3-(4,5-dimethylthiazol-2-yl)-2,5-diphenyltetrazolium bromide) was purchased from Amyjet Scientific (Wuhan, China). The β4 and β1 siRNA kits, as well as EdU Apollo 488 In Vitro Imaging Kit were obtained from Ribobio (Guangzhou, China).

### Cell culture and treatment

HIOECs were cultured in keratinocyte serum-free medium (K-SFM; Gibco) in a humidified atmosphere containing 5% CO2 at 37°C. For confocal or immunofluoresence microscopy, cells were seeded in 24-well plates (1.4×10^5^ cells per well). For real-time PCR, Western blot, and transfection assay, the cells were seeded in 6-well plates (8× 10^5^ cells per well). Cells were given the indicated treatment.

### Integrin function-blocking assays

Cells were preincubated with HYD-1 or functional blocking antibodies against α6 (GoH3), α3 (clone P1B5), β4 (clone 3E1), and β1 (clone 12G10) integrin subunits or mouse control IgG at a final concentration at 20 μg/ml at room temperature for 10 min before KGF treatment.

### Matrix degradation assay

Briefly, 0.2% gelatin in PBS was dropped onto coverslips for 15 min at room temperature. Then, HIOECs were seeded onto gelatin matrix-coated coverslips in 24-well plates and grown for 24 h. An Alexa Fluor^®^ 594 Protein Labeling Kit was used to label gelatin for the extracellular matrix degradation assays. The cells were grown on glass coverslips and given the indicated treatment. Then, cells were washed with PBS, fixed in paraformaldehyde at 37°C for 30 min, and blocked with 10% nonimmune goat serum for 1 h at room temperature. After that, cells were incubated with primary antibody to cortactin at a dilution of 1:100 overnight at 4°C, followed by incubation with Alexa Fluor^®^ 488-conjugated secondary antibody (1:200) and Acti-stain^TM^ 670 phalloidin for 1 h at room temperature. The regions in which 594–gelatin was degraded were analyzed using Photoshop CS6. ImageJ (National Institutes of Health, Bethesda, MD) was used to quantify the percentage of fluorescence reduction in the region under the HIOECs.

### Small interfering RNA (siRNA) transfection

HIOECs at a density of 40% confluence were serum-deprived for 24 h for transfection. Interference of integrin subunits β4 and β1 expression was performed by transfecting Lipofectamine 2000 with integrin-targeted siRNA and the universal negative control siRNA according to the manufacturer’s protocol. Knockdown efficacy was confirmed by Western blot and real-time RT PCR 48 h after transfection.

### Western blot and analysis

The procedure for Western blot was as described in the previous study (Sa et al. 2017). The semiquantitative analysis of Western blots was carried ImageJ software. The expression levels of indicated proteins were revealed by the calculating ratio of a special molecule to GAPDH, α-tubulin or β-Actin.

### Real-Time PCR

The procedure and primer nucleotide sequences for PCR were as described in the previous study (Sa et al. 2017).

### MTT and EdU incorporation assays

HIOECs at 3x10^4^ cells/well were seeded in the 96-well plate and exposed to KGF (10 ng/mL; Proteintech) for 24 h. For MTT assays, the cells were incubated with 20 μL of MTT solution (5 mg/mL) at 37°C for 4 h. The supernatant was discarded, and 150 μL of dimethyl sulfoxide (DMSO) was added to each well. The absorbance at 570 nm was measured with a microplate reader (Bio-Tek). For EdU incorporation assays, the proliferating HIOECs were identified using the Cell-Light EdU Imaging Kit according to the manufacturer’s instruction.

### Statistical analysis

All data were analyzed with Student’s t test at a significance level of *P* < 0.05 in GraphPad Prism 6.0 (GraphPad Software, Inc.). The results are expressed as mean ± SEM for 3 independent experiments.

### Author contributions

G.L. Sa contributed to experiment conception and design; data acquisition, analysis, and interpretation; and drafted the manuscript. Z.K. Liu, J.G. Ren, Q.L. Wan, X.P. Xiong, Z.L. Yu, and H. Chen contributed to data analysis and interpretation; and critically revised the manuscript. Y.F. Zhao and S.G. He contributed to experiment conception, and critically revised the manuscript. All authors declare no potential conflicts of interest for all aspects of the work.

## Acknowledgments

This research was supported by the National Natural Science Foundation of China grant NO. 81371126 to S.G. He, grant NO. 81600385 to J.G. Ren, and grant NO. 81801842 to Z.L. Yu. The authors declare no potential conflicts of interest with respect to the authorship and/or publication of this article.

---

**Supplementary Figure 1.**
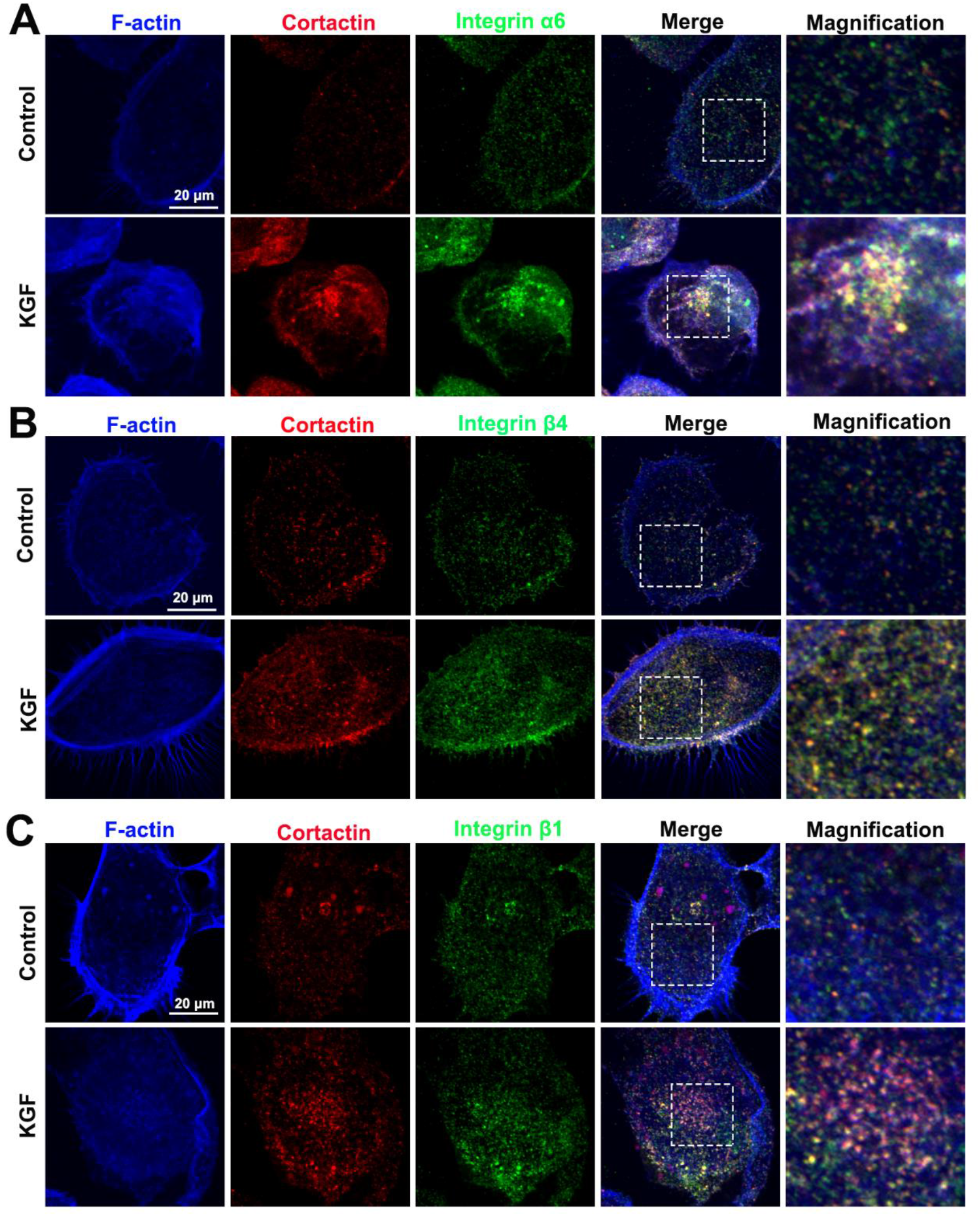
KGF induces the recruitment of podosomal components around F-actin-cortactin complexes in HIOECs. (A) 24 h after KGF treatment, the colocalization among integrin subunit α6 and F-actin-cortactin complexes. (B) The colocalization among integrin subunit β4 and F-actin-cortactin complexes. (C) The colocalization among integrin subunit β1 and F-actin-cortactin complexes. Control: HIOECs cultured in normal K-SFM; KGF: HIOECs cultured in normal K-SFM supplemented with KGF at a concentration of 10 ng/mL.

**Supplementary Figure 2.**
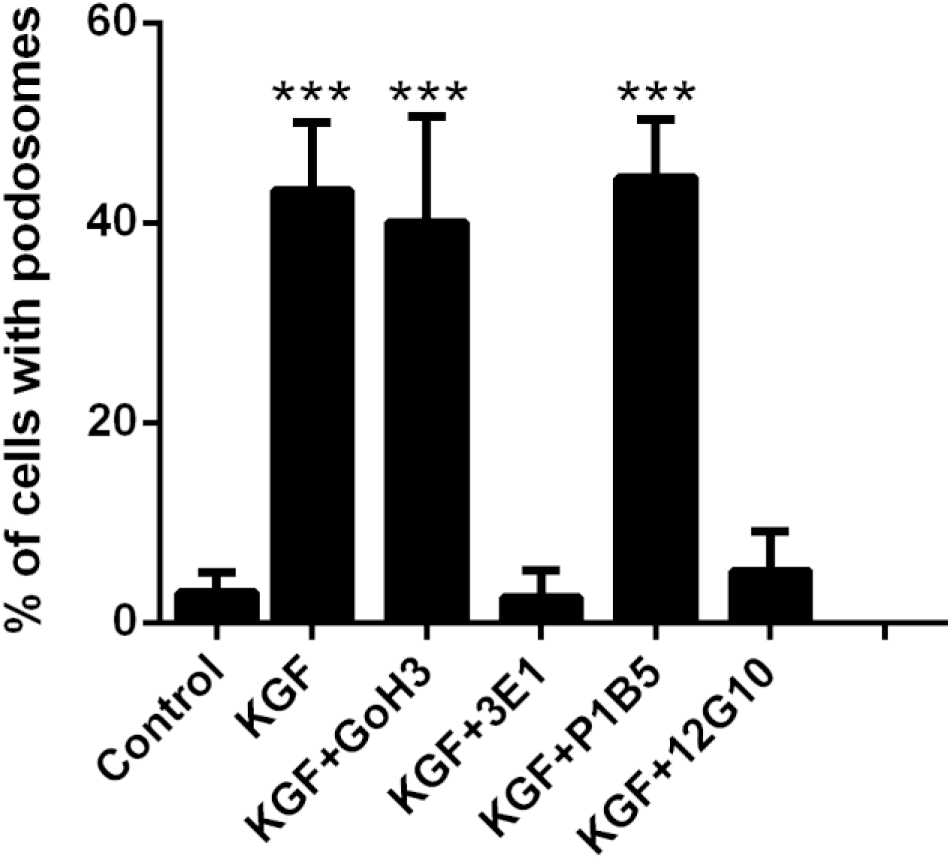
Statistical analysis of the percentage of HIOECs with podosomes in each group. GoH3, function-blocking antibody against integrin subunit α6. 3E1, function-blocking antibody against integrin subunit β4; P1B5, function-blocking antibody against integrin subunit α3; 12G10, function-blocking antibody against integrin subunit β1 . n=3 independent experiments in which 100 cells per experiment were analyzed. Data are expressed as the means ± SEM. \*\**P* < 0.01.

**Supplementary Figure 3.**
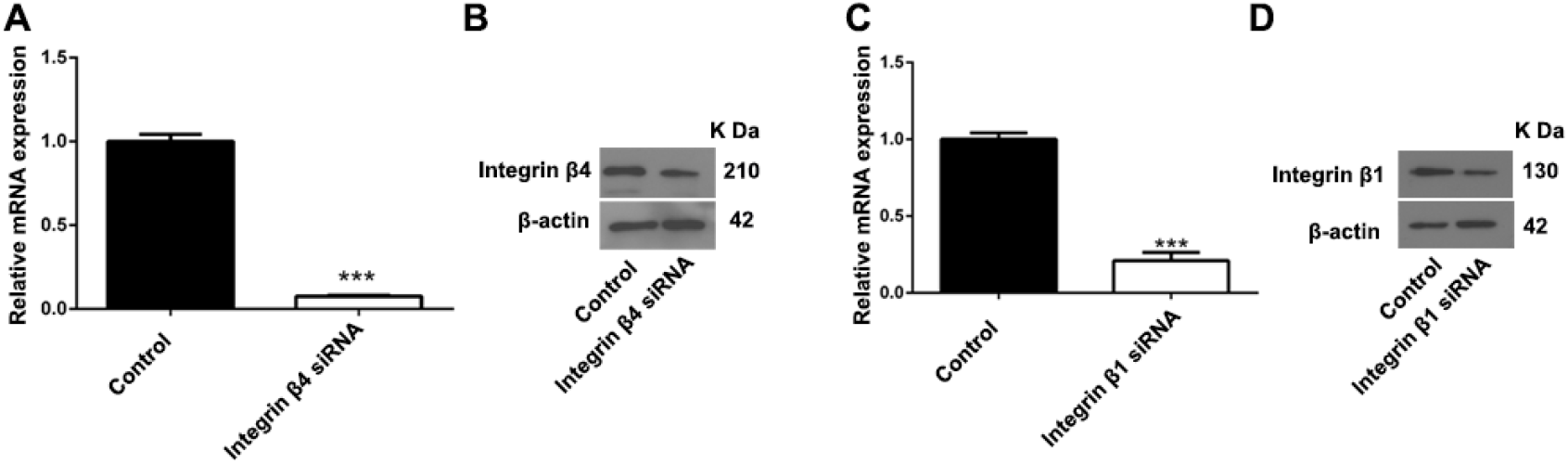
The knockdown efficiency of specific siRNAs against integrin subunits β4 and β1. (a) Real-time polymerase chain reaction analysis of the mRNA expression levels of integrin subunit β4 24 h after incubation with a specific siRNA against integrin subunit β4. (c) Real-time polymerase chain reaction analysis of the mRNA expression levels of integrin subunit β1 24 h after incubation with a specific siRNA against integrin subunit β1. GAPDH was used as a loading control. The experiments were repeated in triplicate. Data are expressed as the mean ± SEM. \*\*\**P* < 0.001. (b) The expression levels of integrin subunit β4 24 h after incubation with a specific siRNA against integrin subunit β4. (d) The expression levels of integrin subunit β1 24 h after incubation with a specific siRNA against integrin subunit β1.

**Supplementary Table 1.**
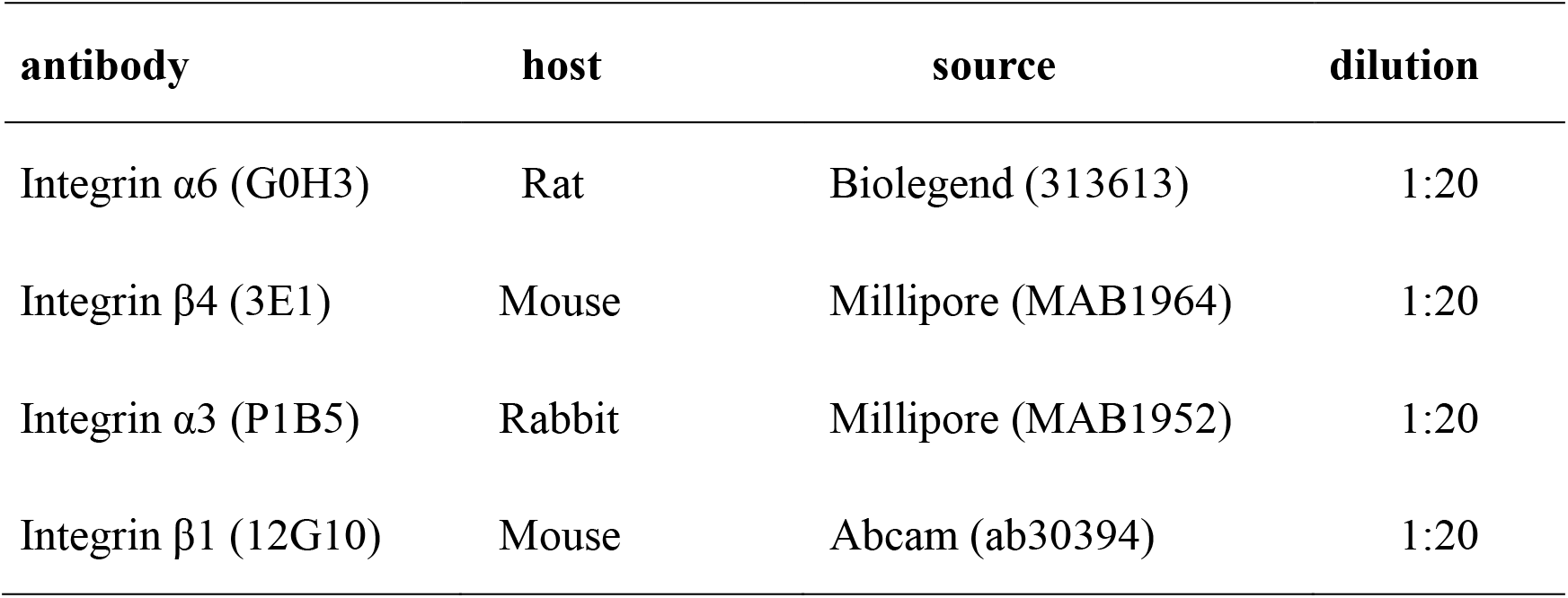
Antibodies used for function blocking assay

**Supplementary Table 2.**
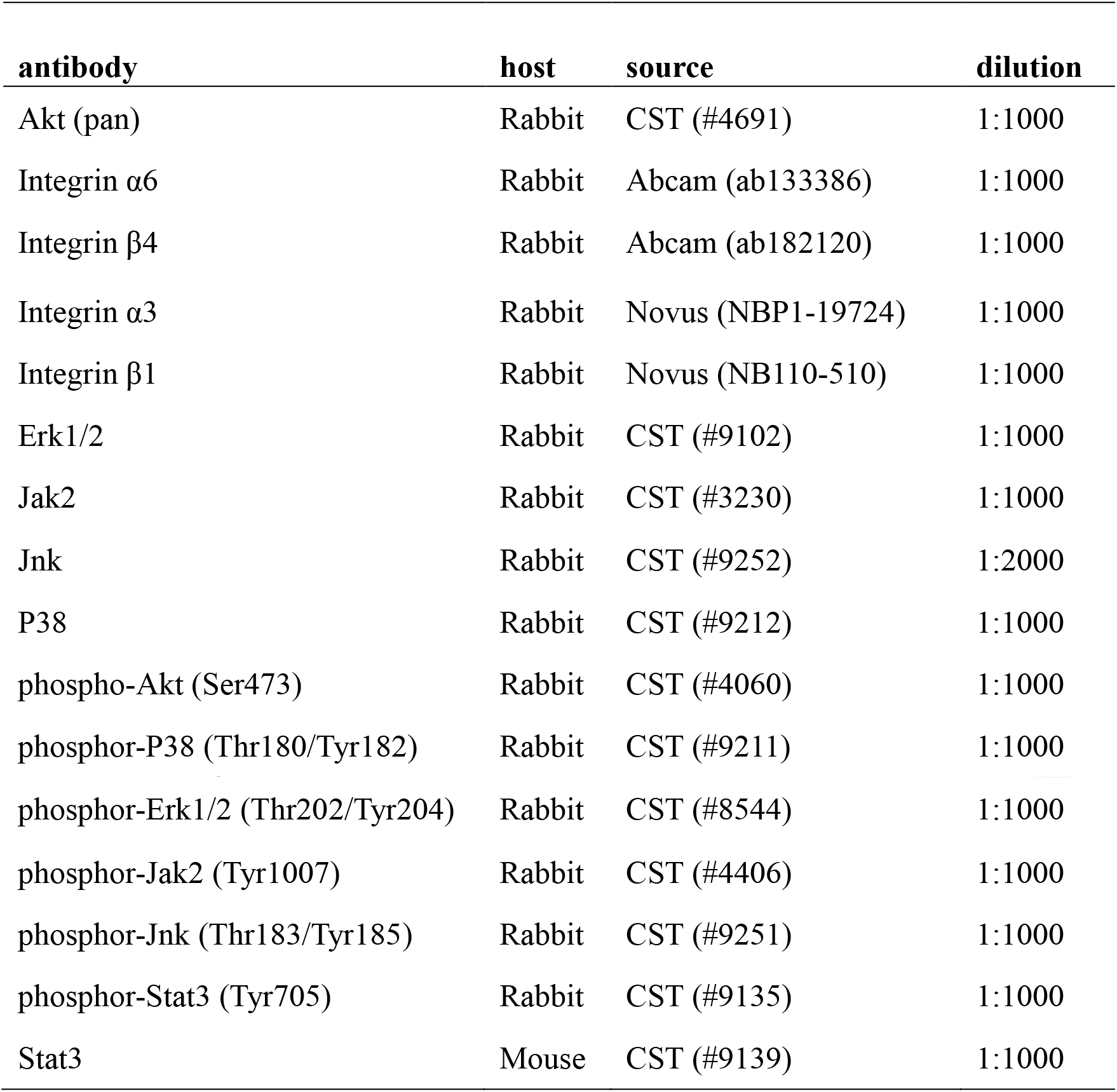
Antibodies used for Western blot

| **antibody** | **host** | **source** | **dilution** |
| --- | --- | --- | --- |
| Akt (pan) | Rabbit | CST (#4691) | 1:1000 |
| Integrin $\alpha 6$ | Rabbit | Abcam (ab133386) | 1:1000 |
| Integrin $\beta 4$ | Rabbit | Abcam (ab182120) | 1:1000 |
| Integrin $\alpha 3$ | Rabbit | Novus (NBP1-19724) | 1:1000 |
| Integrin $\beta 1$ | Rabbit | Novus (NB110-510) | 1:1000 |
| Erk1/2 | Rabbit | CST (#9102) | 1:1000 |
| Jak2 | Rabbit | CST (#3230) | 1:1000 |
| Jnk | Rabbit | CST (#9252) | 1:2000 |
| P38 | Rabbit | CST (#9212) | 1:1000 |
| phospho-Akt (Ser473) | Rabbit | CST (#4060) | 1:1000 |
| phosphor-P38 (Thr180/Tyr182) | Rabbit | CST (#9211) | 1:1000 |
| phosphor-Erk1/2 (Thr202/Tyr204) | Rabbit | CST (#8544) | 1:1000 |
| phosphor-Jak2 (Tyr1007) | Rabbit | CST (#4406) | 1:1000 |
| phosphor-Jnk (Thr183/Tyr185) | Rabbit | CST (#9251) | 1:1000 |
| phosphor-Stat3 (Tyr705) | Rabbit | CST (#9135) | 1:1000 |
| Stat3 | Mouse | CST (#9139) | 1:1000 |

